# Quantification of healthspan in aging mice: Introducing FAMY and GRAIL

**DOI:** 10.1101/2023.11.07.566044

**Authors:** Dudley W. Lamming

## Abstract

The population around the world is graying, and as many of these individuals will spend years suffering from the burdens of age associated diseases, understanding how to increase healthspan, defined as the period of life free from disease and disability, is an urgent priority of geroscience research. The lack of agreed-upon quantitative metrics for measuring healthspan in aging mice has slowed progress in identifying interventions that do not simply increase lifespan, but also healthspan. Here, we define FAMY (Frailty-Adjusted Mouse Years) and GRAIL (Gauging Robust Aging when Increasing Lifespan) as new summary statistics for quantifying healthspan in mice. FAMY integrates lifespan data with longitudinal measurements of a widely utilized clinical frailty index, while GRAIL incorporates these measures and also adds information from widely utilized healthspan assays and the hallmarks of aging. Both metrics are conceptually similar to quality-adjusted life years (QALY), a widely-utilized measure of disease burden in humans, and can be readily calculated from data acquired during longitudinal and cross-sectional studies of mouse aging. We find that interventions generally thought to promote health, including calorie restriction, robustly improve healthspan as measured by FAMY and GRAIL. Finally, we show that the use of GRAIL provides new insights, and identify dietary restriction of protein or isoleucine as interventions that robustly promote healthspan but not longevity in female HET3 mice. We suggest that the routine integration of these measures into studies of aging in mice will allow the identification and development of interventions that promote healthy aging even in the absence of increased lifespan.

## Introduction

As the global population ages, addressing the challenge of age-related diseases has become increasingly urgent. While the field of geroscience has traditionally focused on extending lifespan, there is limited interest in extending the portion of one’s life where an individual is frail, unable to perform activities of daily living, and may be in pain. Instead, the key priority of geroscience should be to increase healthspan, defined as “the period of life spent in good health, free from the chronic diseases and disabilities of aging” (1).

The scientific literature has become replete with discussion of interventions that promote healthspan, increasing exponentially from 0-1 references in PubMed each year from 2000-2004 to 251 references in the year 2022. Many of these references can be found in papers, primarily utilizing mice, stating that various geroprotective interventions increase, improve or extend the healthspan of mice. However, as pointed out by a piece in *Geroscience* as well as in recent online discussions (1), there are no quantitative metrics for measuring healthspan in mice, and thus demonstrating that interventions have a statistically significant benefit to healthspan is not formally possible. This unfortunate gap limits our ability to compare the benefits of different interventions on healthspan – and may lead us to entirely miss interventions that promote healthy aging but do not extend lifespan.

While quantitatively determining healthspan has not been rigorously pursued, two complimentary approaches for assaying healthspan in mice have been developed. The first of these approaches is focused on frailty, defined as an accumulation of multiple deficits and a loss of resilience to adverse events. A frailty index (FI) for mice developed by Dr. Susan Howlett and colleagues a decade ago has become widely utilized as a robust way of measuring frailty in mice that tracks similar clinical frailty indexes in humans (2) and is strongly associated with mortality (3). Calorie restriction (CR), the gold standard for lifespan-extending interventions and the prevention or delay of many age-related diseases, clearly reduces frailty as assessed using the FI (4). An important component of the FI approach is that it is an average of multiple deficits, and thus is robust to the exclusion of specific measurements for logistical or scientific reasons, as well as to the inclusion of additional deficits as they are developed (5). Further, tracking frailty does not generally require specialized equipment and can be performed in a completely non-invasive manner, allowing frailty to be measured repeatedly throughout the lifespan. While these are compelling reasons to measure frailty, there is as yet no standard methodology to integrate longitudinal measurements of FI and lifespan.

The second approach, which has been adopted by a variety of leading aging laboratories, is to assess multiple parameters of healthspan using standardized approaches. A recently published consensus toolbox of protocols to assess healthspan in aging mice includes protocols to assess cardiac function, muscle strength and neuromuscular function, metabolic health, and cognitive function (6). The specific assays selected were chosen based on clinical relevance, reproducibility, a low level of invasiveness or stress for the animal, and a low level of technical difficulty. However, as many of these assays still require injections, blood collection, or anesthesia, and/or involve stress, they will most often be performed only a few times, or perhaps even only once. For these reasons, it may be best to perform these assays on a cohort of mice that will be sacrificed cross-sectionally rather than used for lifespan determination.

Here, we utilize these approaches to define two new quantitative summary statistics to measure healthspan in mice. The first of these, FAMY (Frailty-Adjusted Mouse Years), defines health as a function of frailty, and integrates lifespan data with longitudinal measures of frailty in a matter akin to the widely-used quality-adjusted life year (QALY) in humans (7). The second of these measures, GRAIL (Gauging Robust Aging when Increasing Lifespan), utilizes both lifespan and frailty data but further incorporates information from healthspan assays and the hallmarks of aging. We show that FAMY and GRAIL can be readily calculated from data collected during longitudinal mouse aging studies, and that CR – widely stated in the literature to promote healthspan – increases both FAMY and GRAIL. We further show that the consideration of FAMY and GRAIL can lead to conclusions that are not immediately apparent from the consideration of lifespan, identifying protein restriction as an intervention that increases healthspan without increasing lifespan, highlighting the potential utility of these indexes in identifying interventions that may be able to improve healthspan.

## Results

### The Frailty-Adjusted Mouse Year (FAMY) demonstrates the cost of missing health

In humans, the QALY is the gold standard for measuring the effects, positive or negative, of medical treatments on patients’ lives, and serves as a fundamental component of the cost-effectiveness of medical interventions. 1 QALY is equal to 1 year of life in perfect health. Thus, the total QALY for an individual can be calculated as the area under the curve (AUC) of a graph where quality (utility) of life, a unitless value varying between 0 and 1, is plotted on the y-axis and the survival in years in plotted on the x-axis (7).

We considered that by plotting 1-FI over the lifespan of an animal, we would essentially be able to plot a heathspan curve for each individual mouse in an aging study. When age on the x-axis is plotted in years, we name the AUC of a plot of 1-FI FAMY, for “Frailty-Adjusted Mouse Years” the units of FAMY are “years lived in perfect health” and are the murine equivalent of QALY. In order to “test-drive” the FAMY, we analyzed publicly available lifespan and frailty data from a recent study of C57BL/6J.Nia conducted by experts in the biology of aging and frailty (8). The mice in this study were appropriately long-lived, with a median lifespan of 954 days (**Fig. 1A**), and frailty in this population of mice increased with age as expected (**Fig. 1B**).

**Figure 1.**
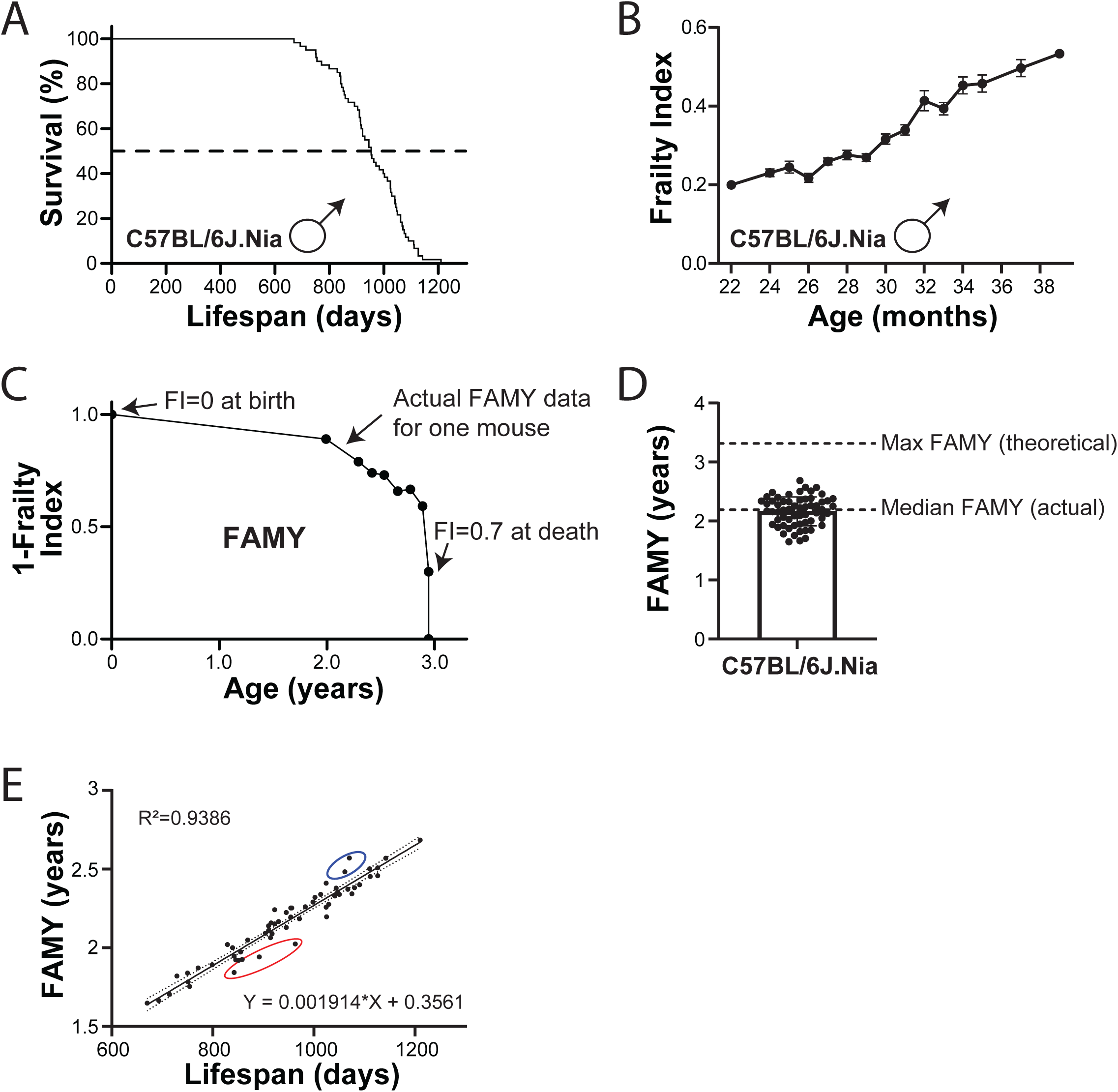
The Frailty-Adjusted Mouse Year (FAMY) as a measure of mouse healthspan. (A-B) Survival (A) and Frailty Index (B) of C57BL/6J.Nia male mice from (8). (C) Illustration of FAMY calculation. The FAMY score for an individual mouse combines the area under the curve of 1-frailty of an individual animal, which varies over time; 1-FI is set to 1.0 at t=0, and to 0.3 at the day of death. (D) Frailty-Adjusted Mouse Years (FAMY) in years was calculated using the survival and frailty data plotted in panels A and B. Max FAMY (theoretical) indicates the FAMY score if the longest-lived mouse in this study remained in perfect health (FI=0) for its entire life. (E) FAMY plotted as function of lifespan. The best-fit linear regression line with 95% confidence bands is shown. (A-B, D-E) n=60 biologically independent mice.

Our calculation of FAMY is performed as explained here and outlined in **Fig. 1C**. We calculated FAMY as the AUC for each individual mouse of a graph of 1-FI, plotting 1-FI from birth until death. In order to plot FAMY throughout the lifespan, we made two simplifying assumptions. First, as aging studies using wild-type mice do not start at birth – many studies start in adult mice and many in mid-life – and frailty assessment may not begin until mid-life or later, we made the simplifying assumption that mice at the beginning of the lifespan study had a FI of 0. Second, we assumed that mice at the end of their life, on the date they are found dead or euthanized due to reaching endpoint criteria, have a FI of 0.7, as the maximum FI score observed for an individual wild-type mouse or human does not exceed 0.7 (3, 9).

We calculated the FAMY for each mouse; the FAMY values primarily grouped between 1.5 and 2.5 years, with a median of 2.19 years (**Fig. 1D**). As expected, FAMY was highly correlated (R^2^=0.9337) with lifespan (**Fig. 1E**). A few points are apparent. First, while the median lifespan of these mice was 2.6 years, the FAMY of 2.19 years reflects that the overall population lost 16.2% of its possible healthspan. Second, the longest-lived mouse at 3.32 years had the highest FAMY, 2.68 years, while the maximum possible FAMY for this mouse was 3.32 years (**Fig. 1D**); thus, this animal lost 19.0% of its possible healthspan to aging. Or, conversely, an intervention that extends healthspan perfectly could increase the healthspan of this mouse by 23.5% without increasing lifespan at all. Finally, while the correlation between FAMY and lifespan is very tight, there is variation, with multiple mice surviving for longer than expected with a poorer healthspan (**Fig. 1E**, red), as well as mice that died sooner than expected with good healthspan (**Fig. 1E**, blue).

### Calorie restriction quantitatively increases healthspan

CR is widely acknowledged as the gold-standard for geroprotective interventions. CR extends the lifespan of a wide range of species, ranging from yeast, rotifer, worms and flies to mice, rats, dogs, and non-human primates, and has shown favorable effects in short-term human trials (10–17). However, the level of CR that optimally extends lifespan may vary by sex and genetic background, and it has been reported that in some strains 40% CR shortens lifespan (18, 19). While a negative effect of CR on the lifespan of certain mouse strains has only been partially replicated (20), these findings have led some to question the safety of dietary interventions like CR particularly as applied to humans in the real world beyond the confines of the laboratory (21).

To begin to address this question, we analyzed the healthspan of mice in a recently completed CR study (22). As shown in **Fig. 2**, the approximately 40% CR regimen utilized here results in significant, 16% increase in the median lifespan of male C57BL/6J mice along with a decrease in frailty as the mice aged (**Fig. 2A-B**). A similar but larger effect on longevity was seen in females, with a 39% increase in median lifespan and a dramatic separation of frailty curves between *ad libitum* (AL) fed controls and mice maintained on a CR diet (**Fig. 2C-2D**).

**Figure 2.**
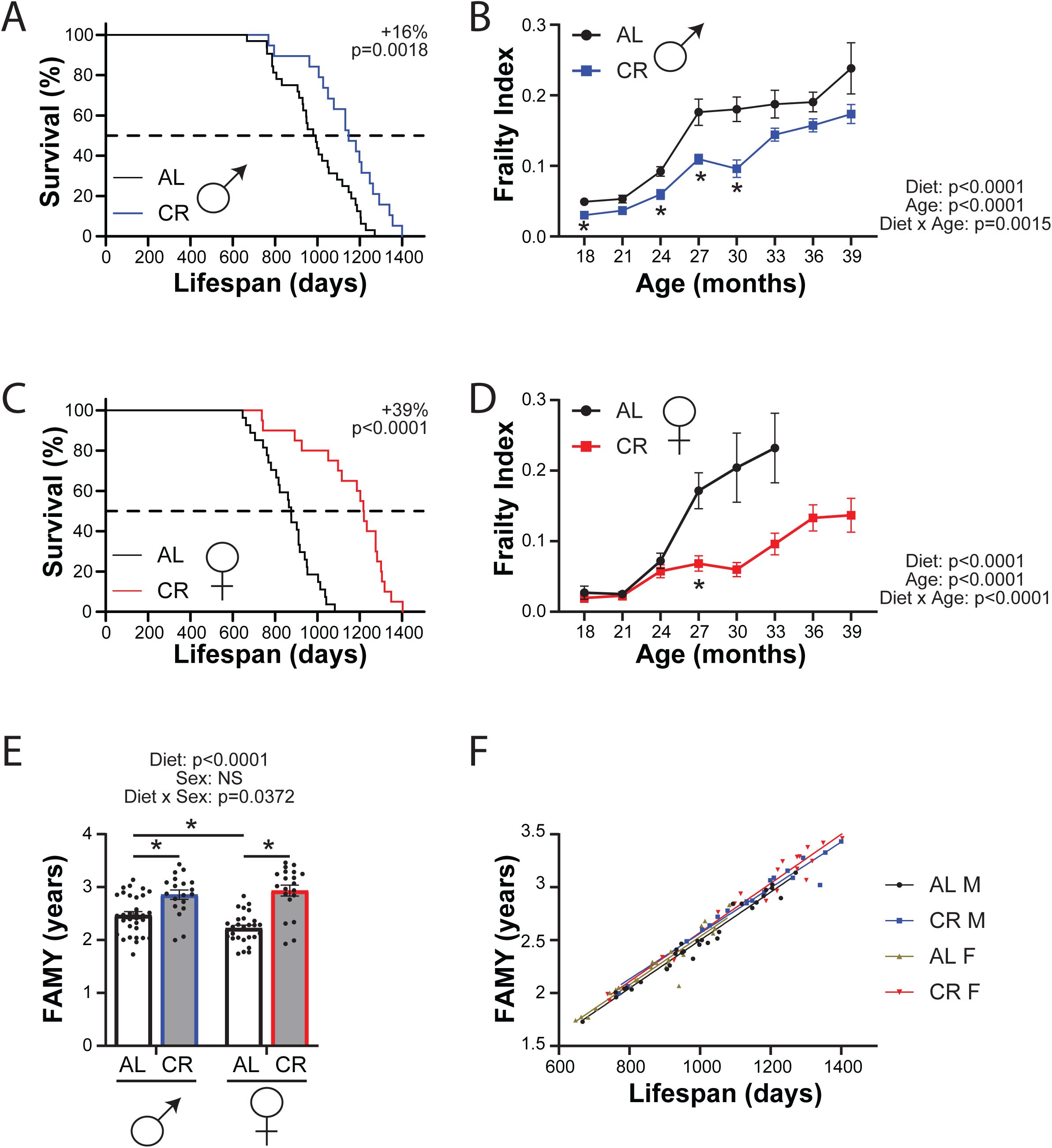
Quantifying the effect of CR on the healthspan of inbred mice with FAMY. (A-B) Survival (A) and Frailty Index (B) of C57BL/6J male mice fed either *ad libitum* or calorie restricted (CR) starting at 10 weeks of age. (C-D) Survival (C) and Frailty Index (D) of C57BL/6J female mice fed either *ad libitum* or calorie restricted (CR) starting at 10 weeks of age. (E) FAMY calculated using the survival and frailty data plotted in panels A-D. (F) FAMY plotted as function of lifespan. (A, C) Log-rank test for AL vs. CR. (B, D) Mixed-effects model (REML) for time and diet with post-hoc Sidak’s test, *p<0.05. (E) Sidak test following 2-way ANOVA. \**p*<0.05, P-values for the overall effect of Diet, Sex and the interaction represent the significant p-values from the two-way ANOVA. (F) The best-fit linear regression line for each group is shown. (A-F) n=32 AL males, 19 CR males, 27 AL females, 20 CR females. Data represented as mean ± SEM.

As above, we calculated healthspan using FAMY. In accordance with the general belief that CR is a healthspan-extending intervention, we find that CR significantly increases FAMY in both male and female mice (**Fig. 2E**). Interestingly, while the average healthspan of male mice is increased 15.6% by CR, similar to the effect on lifespan, the healthspan of female mice increases only 31.9%, less than the effect of CR on lifespan. To get a better idea of whether this represented a real difference in the effect of CR on male and female healthspan, we examined the correlation between FAMY and lifespan for each group of mice (**Fig. 2F**). As we expected, the correlation between FAMY and lifespan for each group was quite high, with an R of 0.9 or greater.

We plotted regression lines for each group, and we noted that while there was no statistically significant difference in slopes or intercepts between any of the groups, the CR groups were consistently higher than the AL fed groups. As there was no effect of sex, we pooled the sexes, and found that the y-intercept of CR-fed mice was 0.0297 healthy years greater than that of AL-fed mice (p=0.004). Using the regression equations for AL and CR mice, we calculated that an AL-fed mouse that died after 2.5 years would have an average FAMY of 2.32 years, while a CR-fed mouse that lived 2.5 years would have an average FAMY of 2.37 years – an increase of only 2.3%. Thus, it seems that at least in this particular experiment, CR generally increased healthspan primarily as a result of the longer lifespan; a CR-fed mouse that died at the same age as an AL-fed mice would be only marginally healthier.

### Calorie restriction and CR-mimietic interventions quantitatively increase healthspan in independent data sets

In order to examine the broad utility of FAMY as an approach to healthspan interventions, we decided to examine FAMY in independent datasets generated by other laboratories. This was somewhat difficult, as several studies we examined collected healthspan data either at single time points or from only a subset of animals whose lifespans were also determined. Fortuitously, two recent studies collected longitudinal frailty data and determined the lifespan of genetically heterogeneous Diversity Outbred (DO) mice on one of several different diet regimens (23, 24). In these studies, mice were placed on *ad libitum* (AL) diet, intermittently fasted (1D and 2D represent groups that fasted once or two consecutive days per week in the first study, respectively, while IF1 and IF2 represent mice that were fasted either one or two days per week in the second study, respectively), or placed on either a 20% or 40% CR diet (CR20 and CR40, respectively). The primary difference between the studies is that diets were initiated at 6 months of age in the first study (24), and initiated at 2.5 years of age in the second study (23).

Surprisingly based on previous literature suggesting CR works poorly when initiated late in life (25), both studies found that mice placed on either 20% or 40% CR lived significantly longer, while intermittent fasting did not significantly extend lifespan (23, 24). Calculating healthspan from the longitudinal frailty data, we found that 20% and 40% CR, but not intermittent fasting, significantly extended healthspan (**Fig. 3A-B**). In agreement with the idea that CR is more potent in younger animals, mice that started on 40% CR at 6 months of age had a 25.6% increase in healthspan, while mice starting 40% CR at 2.5 years of age had a 17.8% increase in healthspan relative to AL-fed mice.

**Figure 3.**
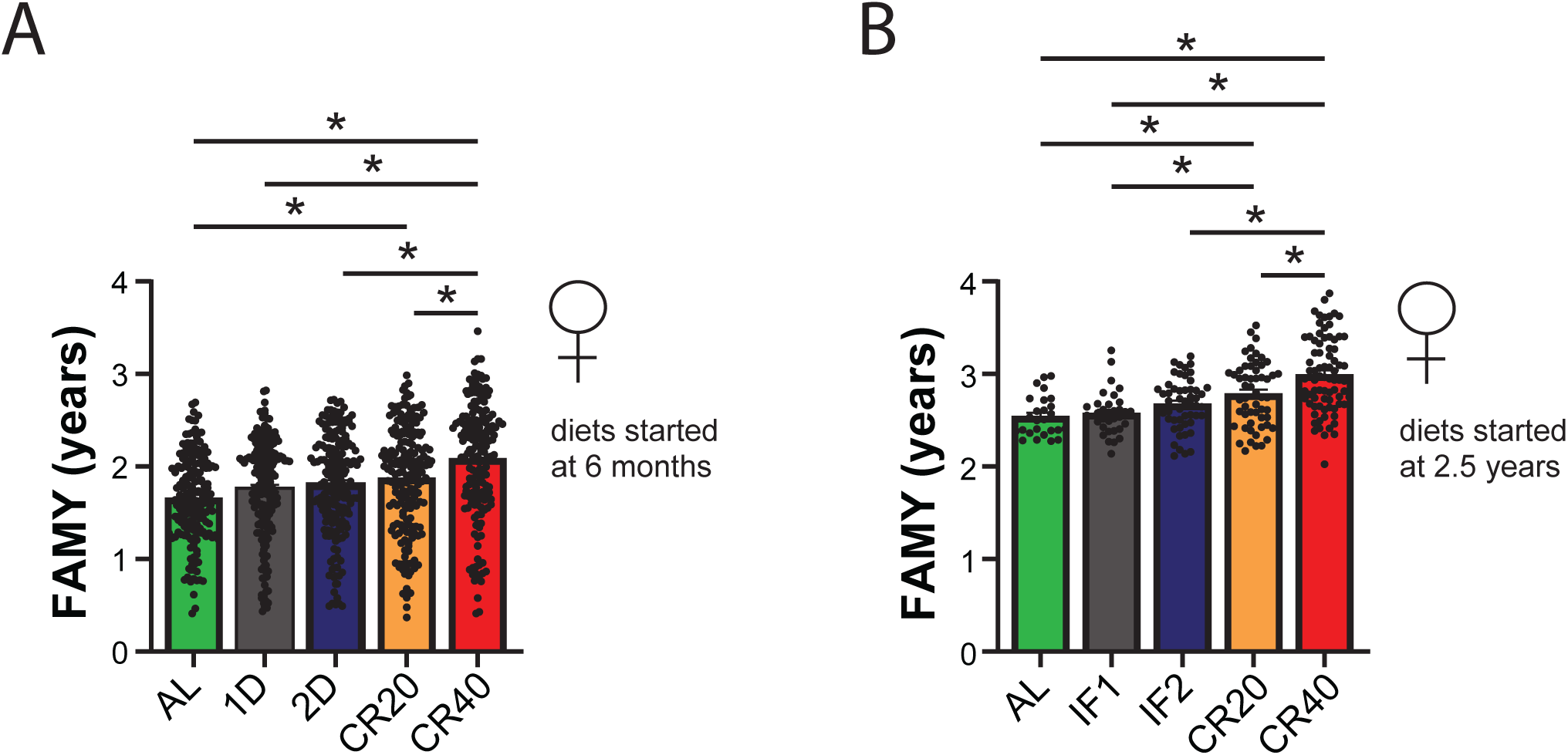
CR, but not fasting, improves the healthspan of Diversity Outbred mice. (A-B) Survival and healthspan (FAMY) of DO mice fed the indicated diets from 2.5 years of age were plotted and calculated using the surivial and frailty data from Luciano *et al.*, 2024 (23). (C) Healthspan (FAMY) of DO mice fed the indicated diets from 6 months of age was calculated during the surivial and frailty data from Di Francesco *et al.*, 2023 (24).

### Restriction of isoleucine quantitatively increases male healthspan as assessed by FAMY

There is a growing awareness among both scientists and the public that a calorie is not “just a calorie,” and that calories from different macronutrient sources may have effects on metabolism and healthy aging that are not simply related to their energy content (26). For example, restriction of dietary protein or of methionine extends the lifespan of rodents (27–30), and lifelong restriction of the three branched-chain amino acids (BCAAs; leucine, isoleucine, and valine) extends lifespan and reduces frailty in male C57BL/6J mice (31).

Building on our recent discovery that restriction of isoleucine alone is sufficient to improve metabolic health and is necessary for the benefits of protein restriction in C57BL/6J male mice (32), we decided to test the effects of isoleucine restriction (IleR) on healthy aging. Initiating IleR in 6-month-old genetically heterogeneous mice of both sexes, we recently found that IleR had profound effects on longevity vs. an amino acid-defined Control diet, extending median lifespan of male by 33% and females by 6%, and extending the maximum lifespan of male mice (**Figs. 4A, B** and (33)). In contrast, neither male nor female mice fed an amino acid defined protein restricted diet (PR) in parallel lived longer than Control-fed mice. Male mice fed the PR or IleR diets had reduced frailty vs. Control-fed at multiple time points starting at 24 months of age, while female PR and IleR-fed mice had reduced frailty only at 24 months of age (**Figs. 4C, D**).

**Figure 4.**
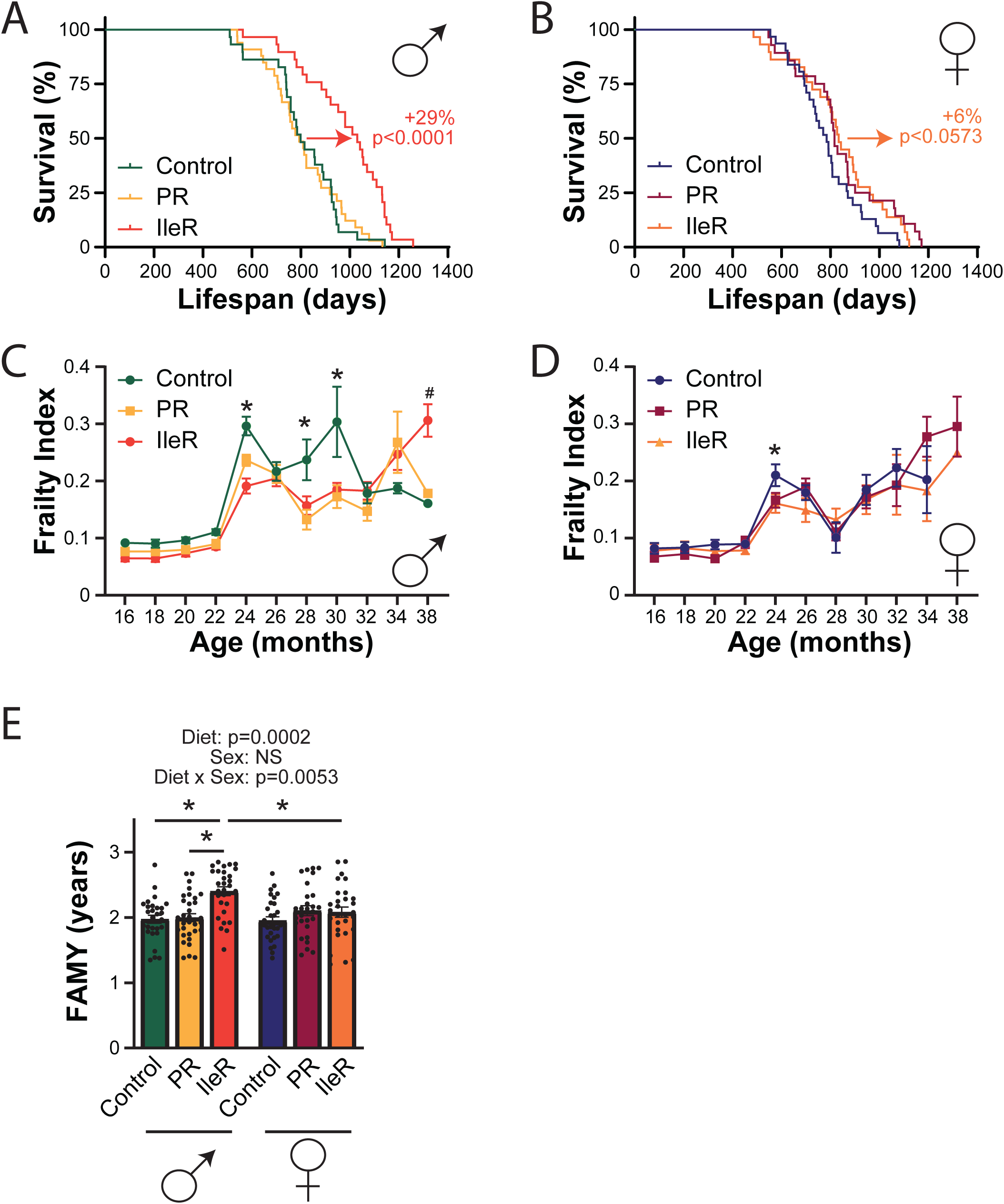
Quantifying the effect of reduced protein or isoleucine on the healthspan of genetically heterogeneous mice with FAMY. (A-B) Survival of male (A) and female (B) UM-HET3 mice fed the indicated diets starting at 6 months of age. (C-D) Frailty index of the mice plotted in panels A-B. (E) FAMY calculated using the survival and frailty data plotted in panels A-D. (A-B) Log-rank test for Control vs. IleR. (C-D) Mixed-effects model (REML) for time and diet with post-hoc Tukey’s test, *p<0.05 Control vs. PR and Control vs. IleR; #p<0.05 Control vs. IleR only. (E) Tukey test following 2-way ANOVA. \**p*<0.05, P-values for the overall effect of Diet, Sex and the interaction represent the significant p-values from the two-way ANOVA. (A-E) n=29 Control male, 33 PR male, 29 IleR male; 31 Control female, 28 PR female, 29 IleR female. Data represented as mean ± SEM.

To assess the effects of these diets on healthspan, we calculated the FAMY for each mouse in the lifespan study. We observed an overall effect of diet, as well as a sex-diet interaction, on FAMY. IleR increased FAMY in males, but did not significantly increase FAMY in IleR-fed females (**Fig. 3E**). The effect of IleR on median FAMY (+24%) was lower than the effect of IleR on median lifespan (29%), suggesting that as we observed with CR, IleR increases healthspan in males slightly less than it increases lifespan.

### Integrating measures of healthspan and the hallmarks of aging

A key advantage of frailty measurements is that most of the measures included in the clinical frailty index for mice are non-invasive (exceptions include rotarod performance and rectal temperature, and we omit these from longitudinal frailty assessment in our laboratory). However, there are many other procedures to assess the health of aging mice. For example, a toolbox recently published by the MouseAGE international consortium includes detailed procedures for common assays of metabolic health, physical and cognitive performance, and cardiovascular function (6). Twelve hallmarks of aging were recently described; assays to examine the hallmarks of aging often require blood or tissue samples, typically from euthanized mice (34). While these healthspan assays and the hallmarks of aging provide valuable information about the health of mice, for scientific and logistical reasons they cannot be performed frequently across the lifespan. We thus decided to construct a new measure, the GRAIL (Gauging Robust Aging when Increasing Lifespan) to integrate information about both frailty and the health/hallmarks.

We decided to calculate GRAIL as the sum of two components as outlined in **Figure 5A**. Similar to the FAMY, we calculated the GRAIL frailty component (GRAIL-FI) by determining the AUC for each individual mouse of a graph of 1-FI, with the FI on the day of death set equal to 0.7. Unlike for FAMY, we do not include the AUC between 0 and 0.3 of 1-FI; thus, GRAIL-FI closely approximates 70% of FAMY.

**Figure 5.**
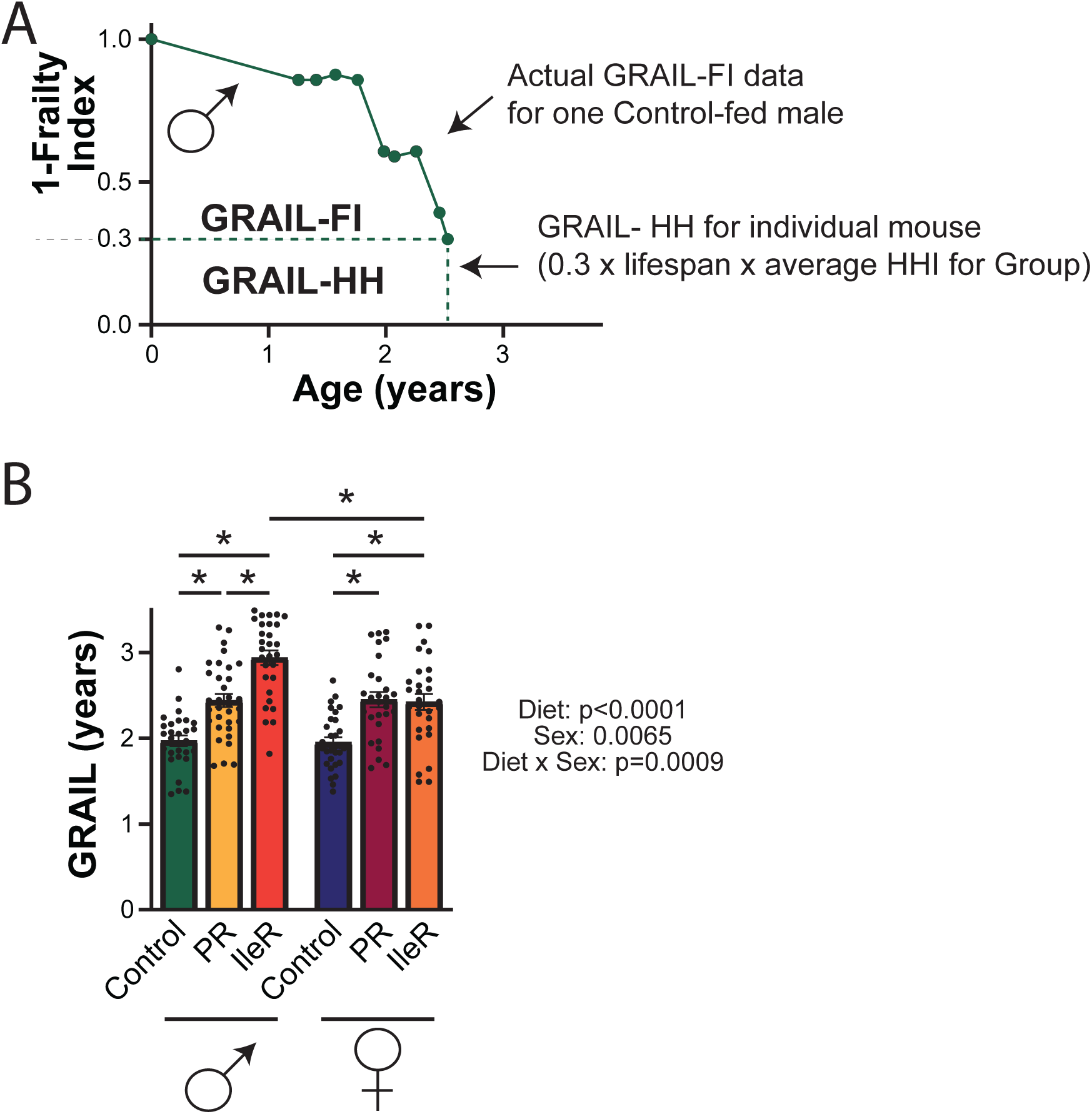
Integrating frailty, healthspan assays, and the hallmarks of aging with GRAIL (Gauging Robust Aging when Increasing Lifespan). (A) Illustration of GRAIL calculation. The GRAIL score for an individual mouse combines the area under the curve of 1-frailty of an individual animal (GRAIL-FI), which varies over time, and information about the average effect on healthspan assays and the hallmarks of aging (GRAIL-HH). (B) GRAIL calculated using the survival and frailty data plotted in Fig. 3 **A-D**. Tukey test following 2-way ANOVA. \**p*<0.05, P-values for the overall effect of Diet, Sex and the interaction represent the significant p-values from the two-way ANOVA. n=29 Control male, 33 PR male, 29 IleR male; 31 Control female, 28 PR female, 29 IleR female. Data represented as mean ± SEM.

The GRAIL health/hallmarks component (GRAIL-HH) was calculated not for individual animals but for specific groups of animals that are being compared (e.g., WT CR males in **Fig. 2**). We determine a health and hallmarks index (HHI) as the average of the list of 22 items in **Table 1**. If the value of HHI is comparable to that value for the wild-type untreated control group, we will score the value as 1.0, with 0 indicating a statistically significant impairment in the HHI component relative to the control group, and 2.0 indicating a statistically significant improvement relative to the control group.

In the event that impairment or improvement in an item is for only a portion of the lifespan or is considered only mild, 0.5 or 1.5 may be scored, respectively. Items for which no information is available – for example, telomere length was not examined in the PR/IleR study discussed in Figure 3 – are simply omitted. Finally, GRAIL-HH for an individual mouse is calculated by multiplying the average HHI across all scored items by both 0.3 and the lifespan of the lifespan of the animal. **Figure 5A** illustrates the area of GRAIL-HH for an animal in the control group with a HHI of 1.

**Table 1:**
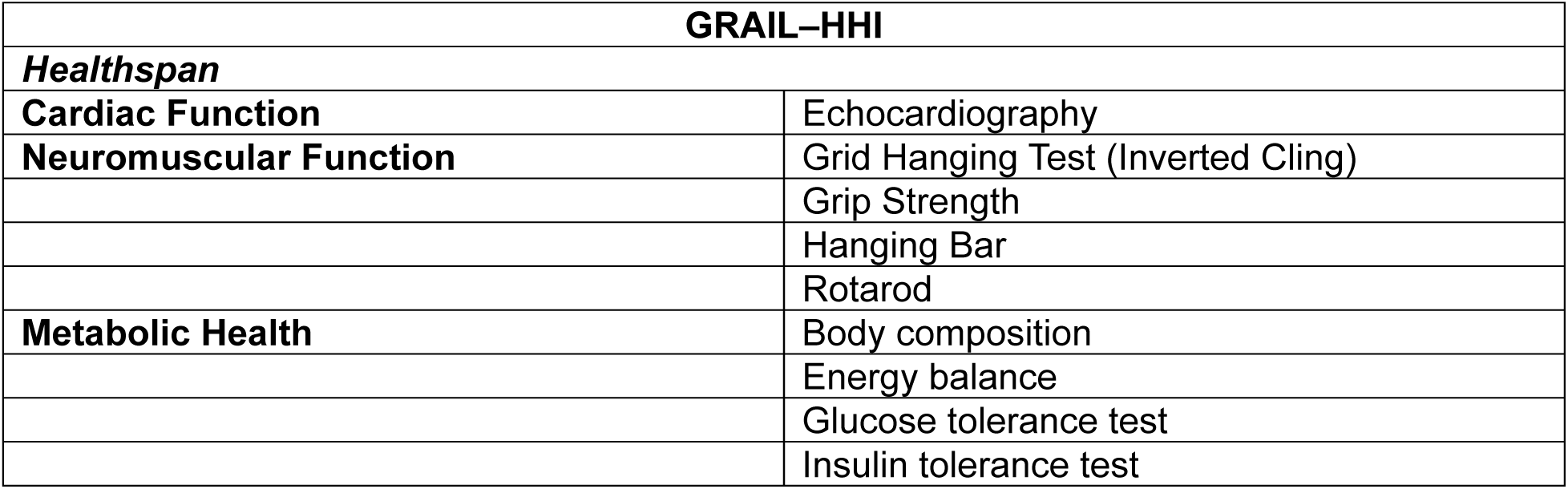
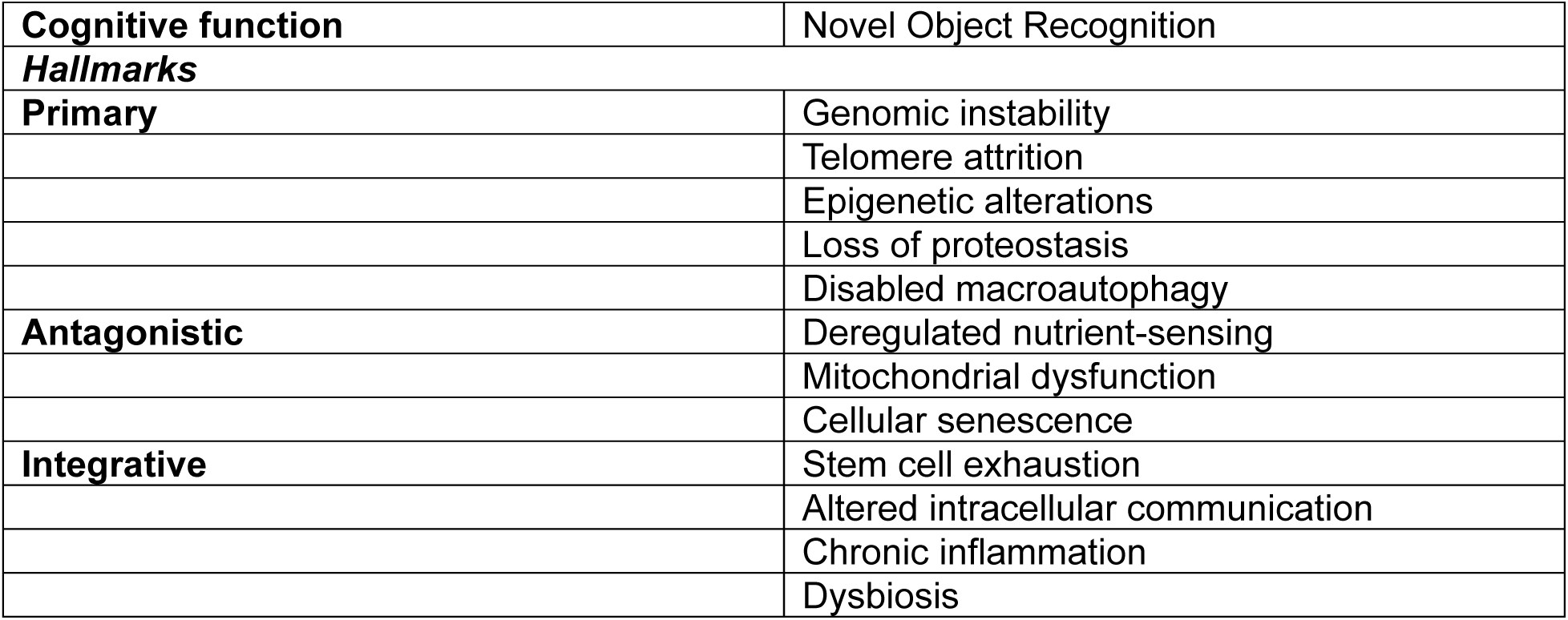
Twenty-two Items that may be scored in the GRAIL health/hallmarks (GRAIL-HH) component.

| GRAIL-HHI |  |
| --- | --- |
| **Healthspan** |  |
| **Cardiac Function** | Echocardiography |
| **Neuromuscular Function** | Grid Hanging Test (Inverted Cling) |
|  | Grip Strength |
|  | Hanging Bar |
|  | Rotarod |
| **Metabolic Health** | Body composition |
|  | Energy balance |
|  | Glucose tolerance test |
|  | Insulin tolerance test |
| **Cognitive function** | Novel Object Recognition |
| **Hallmarks** |  |
| **Primary** | Genomic instability |
|  | Telomere attrition |
|  | Epigenetic alterations |
|  | Loss of proteostasis |
|  | Disabled macroautophagy |
| **Antagonistic** | Deregulated nutrient-sensing |
|  | Mitochondrial dysfunction |
|  | Cellular senescence |
| **Integrative** | Stem cell exhaustion |
|  | Altered intracellular communication |
|  | Chronic inflammation |
|  | Dysbiosis |

We calculated the GRAIL for HET3 mice fed either PR or IleR diets starting at 6 months of age, determining the GRAIL-FI for each individual animal and then adding to it the GRAIL-HH for each animal on the basis of group assignment (**Fig. 5B**). Similarly to the FAMY, we determined that there was a significant effect of diet and a diet-sex interaction; however, we also observed a significant effect of sex. A notable difference between FAMY and GRAIL is that that we observe that IleR increases healthspan not just in males, but in females; and that PR – despite a minimal effect on lifespan – improves healthspan in both sexes. These findings agree generally with our observation of reduced frailty in PR-fed males, and the improved response on both PR and IleR-fed animals on healthspan assays. In further agreement, we observed the greatest healthspan in male IleR-fed mice. Notably, IleR increased median healthspan 52% in males as compared to a 29% increase in median male lifespan; a similar effect, with IleR increasing median healthspan by 28% vs. a 6% increase in lifespan, was observed in females. These results suggest that rather than simply extending lifespan, IleR adds “life to years” by compressing the fracation of life spent in poor health.

## Discussion

While it is widely stated that CR and other geroprotective interventions improve healthspan, the effect of these diets and drugs has not been quantitatively or statistically demonstrated due to lack of integrative healthspan metrics. This has likely slowed progress in geroscience, as the lack of quantitative metrics limits our ability to compare the effect of different interventions on healthspan. Further, it will allow us to settle long-standing debates about the effect of widely studied interventions including CR and rapamycin on the healthspan of mice (18, 19, 35, 36). Finally, the use of FAMY and GRAIL may permit us to use advanced techniques, such as genetic mapping, to identify quantitative trait loci and specific gene alleles that contribute to variation in the effect of interventions on not just lifespan, but healthspan, between different strains and sexes of mice.

Here, we have established FAMY and GRAIL as new quantitative summary statistics of healthspan. We have demonstrated that at least one specific CR regimen in C57BL/6J mice extends not only lifespan, but also healthspan as measured by FAMY, in both males and females. Intriguingly, the increase in healthspan by CR is primarlily the results of most CR-fed mice living longer than AL-fed mice; AL and CR-fed mice that happen to die at approximately the same age have similar healthspans.

We have also shown that an IleR diet increases both lifespan and healthspan in male HET3 mice as assessed using both FAMY and GRAIL. Further, use of GRAIL reveals that both IleR and PR diets extend healthspan, despite the small effects of IleR on lifespan in females and the lack of a longevity effect of PR in either sex. Importantly, we see that IleR adds healthspan in excess of its effects on lifespan.These results not only demonstrate that PR can uncouple healthspan from lifespan, but shows that these summary statistics can help identify interventions that improve healthspan and add healthy years of life, despite not increasing median lifespan.

FAMY and GRAIL provide insight into the cost of frailty to healthy aging. Here, we find that frailty shortens the healthspan of mice by approximately 16-20% from the maximum possible healthspan given a fixed maximum lifespan. This percentage of time is similar to the concept of the “lost decade” in human health – the idea that people typically lose 10-20 years of high quality life due to disability and frailty (37). While the NIA Interventions Testing Program (ITP) as well as other groups have over the past two decades have identified many agents that extend lifespan, going forward tracking healthspan may allow the identification of agents that are just as effective (or more so) at adding healthy years to life. Once such intervention might be metformin, which did not extend lifespan in HET3 mice when tested by the ITP, but has been shown to extend healthspan in assays by others (38, 39).

The FAMY and GRAIL healthspan indexes described here have limitations that may need to be designed around in future versions of these statistics (e.g., FAMY2 or GRAIL2). First, for both FAMY and GRAIL-FI, we need longitudinal frailty data from across the lifespan, which may not always be available. The calculation here assumes that frailty increases linerarly between birth and the first time point measured; while this is likely a good assumption when studies begin when frailty is still low (e.g. at 18 months of age), when studies begin late in life – for example as in **Fig. 3B** when studies begin at 30 months of age – this is not likely true, and leads to discrepancies (see difference in FAMY between **Fig. 3A** and **Fig. 3B**). Calculating change in FAMY between groups, starting not from birth but from the start of the intervention, might better reflect the true impact of the intervention on remaining healthspan.

Importantly, while occasional missing frailty values could potentially be accounted for using imputation (40), mouse values for both lifespan and frailty after a certain age are both missing for mice that are censored for log-rank analysis – for example, because they were sacrificed for cross-sectional examination of tissues or molecular signaling – and thus cannot be used to calculate either FAMY or GRAIL-FI. Secondly, while we have utilized a subset of the 31 original frailty items (2), removing specific items such as body composition that we examine in healthspan assays, these and additional frailty measures could be incorporated here as well. These could include recently developed laboratory measure of frailty, including blood pressure, blood chemistry, and inflammatory markers (5).

Similarly, additional healthspan assays or hallmarks of aging could readily be incorporated into GRAIL. The individual tests utilized in GRAIL could also potentially be weighted differently. Finally, investigators will need to make principled and justifiable choices – ideally before commencement of the study via a standardized protocol – regarding cutoffs regarding the assessment of individual GRAIL-HH assessments, and how to score assessments if different assays provide discordant data (for instance, if two different measures of inflammation, of varying sensitivity, provided different values). Despite these and other limitations, 90% of respondents to a survey conducted at the 2023 Masoro-Barshop Conference on Aging following a presentation about these new summary statistics agreed with the statement that “we can quantitatively measure healthspan using FAMY and/or GRAIL.”

In conclusion, we have developed two new summary statistics, FAMY and GRAIL, for the quantification of healthspan in studies of aging mice. These metrics allow the integration of lifespan and frailty data (FAMY) as well as information from healthspan assays and the hallmarks of aging (GRAIL). We have utilized these new metrics to show that CR promotes healthspan in C57BL/6J mice of both sexes, largely due to the benefits to longevity. Further, PR and IleR can promote not only lifespan, but also healthspan; and that IleR in particular can add healthy years of life in excess of its effects on lifespan, while PR shows that improvements in healthspan can be uncoupled from increased longevity. Additional research remains to be conducted to determine the quantitative effects of other geroprotective interventions, and to identify previously unrecognized geroprotective interventions that improve healthspan without extending lifespan. While additional research will be required, and future versions of FAMY or GRAIL may enhance our ability to quantitatively assess healthspan, our results suggest that FAMY and GRAIL will be widely useful in quantifying the effects of interventions on healthspan.

## Materials and methods

The lifespan and frailty data analyzed here were taken from the supporting information files accompanying the original publications analyzed here: Yu *et al.*, 2019, *Cell Reports*; Schultz *et al.*, 2020, *Nature Communications*; and Green *et al.*, 2023, *Cell Metabolism*. Data was analyzed using Excel, Prism Graphpad, and R.

The techniques used to calculate FAMY and GRAIL are described briefly in the Results section and are detailed here. Sample code and formatted input and output data for the calculation of FAMY and GRAIL.fi is provided as Sample Data In, Sample Data Out, and Sample Code 1.

To calculate FAMY:

1. Collect longitudinal frailty index (FI) data for each individual animal in a lifespan study.
2. Calculate 1-FI at each age (in days of age) where you have FI data.
3. Add three additional datapoints for each mouse:

a. 1-FI = 0 at 0 days of age;
b. 1-FI = 0.3 at the date of death in days;
c. 1-FI = 0 at the date of death in days.
4. Plot 1-FI as a function of age in days
5. Determine the AUC for each mouse
6. Divide by 365 to convert units to years

GRAIL:

1. Calculate GRAIL-FI for each mouse by calculating FAMY as above, and then subtracting (0.3 x age at death in years).
2. Calculate GRAIL-HH as follows:

a. For each grouping of mice (e.g., Low AA-fed females in Figs. 3 and 4), score each of the 22 items listed in Table 1.

i. Enter N/A for items that were not assessed
ii. Enter 1.0 if the health of the group on that item (e.g., glucose tolerance) is comparable to the value of an untreated wild-type control group;
iii. Enter 0 if the assessment of the group on that item is impaired relative to the untreated wild-type control group (e.g., glucose tolerance AUC is increased);
iv. Enter 2.0 if the assessment of the group on that item is improved relative to the untreated wild-type control group (e.g., glucose tolerance AUC is decreased).
v. Modifications: a) In the event of impairment or improvement for only a portion of the lifespan, 0.5 or 1.5 may be scored, respectively. b) values which reflect the degree of improvement could be utilized, customized for each individual item. E.g., 1.5 could reflect a 0-30% improvement in glucose tolerance or a 0-5% reduction in adiposity.
b. Average all assessed items, and then multiple this average by (0.3 x age at death in years).
3. Calculate GRAIL for each mouse by adding the individual GRAIL-FI score to the group GRAIL-HH score.

## Supporting information

Sample Data In

Sample Data Out

Sample Code R

## Data availability

All underlying data has been published as supporting information for Yu *et al.*, 2019, *Cell Reports*; Schultz *et al.*, 2020, *Nature Communications*; Green *et al.*, 2023, *Cell Metabolism;* Luciano *et al.*, 2024, *bioRxiv;* and Di Francesco *et al.*, 2023; *bioRxiv*. Results presented here vary slightly from those presented in Green *et al.*, 2023, *Cell Metabolism* as censored data (e.g. frailty and survival data from mice sacrificed cross-sectionally for molecular analysis) included in the original publication are excluded here as these animals are not included in the calculation of FAMY or GRAIL. Source data for the figures is provided in Source Data 1.

## Code availability

Sample code and formatted data for the calculation of FAMY and GRAIL.fi is provided as Sample Data In, Sample Code 1, and Sample Data Out. Note that the values calculated are in units of days, and must be divided by 365 to calculate FAMY and GRAIL.fi in years.

## Acknowledgements

We would like to thank members of the Lamming laboratory, the organizers and participants at the 2023 Masoro-Barshop Conference on Aging, and Dr. Joseph Baur for their comments and feedback, Dr. Dena E. Cohen for help with data analysis, and Dr. Gary Churchill and colleagues for providing data for their pre-printed studies. The Lamming lab is supported in part by the NIA (AG056771, AG062328, AG081482, and AG084156), the NIDDK (DK125859), and startup funds from UW-Madison. The content is solely the responsibility of the author and does not necessarily represent the official views of the NIH.

