## Supplementary material for "Quantification of healthspan in aging mice: Introducing FAMY and GRAIL": Sample Code R

#####################Code for new FAMY calculations#################################

#This code assumes an input data file meeting the following specifications:
#1. Input data is in "long" form, 1 row per animal per time point
#2. Input variables are defined as follows:
#id = character variable, Unique animal identifier
#doa = numeric variable, Days of age at the time of measurement 
#fi = numeric variable, frailty index (artificially set to 0 at doa = 0 and 1 upon final day of lifespan; 
#     this is then reset to 0.7 on final day of lifespan by the code)
#3. Output data consists of one row per animal
#4. Output variables are defined as follows:
#id = character variable, Unique animal identifier
#famy = numeric variable, calculated value of FAMY using final fi of 0.7
#grail.fi = numeric variable, calculated value of FAMY - 0.3 * lifespan

#clear environment
rm(list=ls())

#load required packages
library(dplyr)
library(pracma)

#load starting data, assumes this is stored locally as a .csv file
in.path <- "C:/Users/dlamming/Documents/Manuscripts in Preparation/QAMY paper/edited for second submission to Nature aging/Sample Data.csv" #replace with the location of the desired file
input <- read.csv(in.path)
input <- input %>% filter(is.na(fi)==FALSE)

#reset final value of FI to 0.7
#calculate fi.1 as 1 - fi for each data point
input$fi <- ifelse(input$fi == 1, 0.7, input$fi)
input$fi.1 <- 1 - input$fi

#sort the dataframe to ensure that observations are in chronological order within each animal
input <- input %>% arrange(id, doa)

#calculate AUC for each animal and output the result as a new data frame

#create vector of unique values of id 
id.vec <- input %>% distinct(id)
id.vec <- id.vec$id

#blank vector to hold results
famy.vec <- vector(mode = "double", length = length(id.vec))

for (i in 1:length(id.vec)){
  
  mouse <- id.vec[i]
  
  set <- input %>% filter(id == mouse)
  
  x <- as.vector(set$doa)
  y <- as.vector(set$fi.1)
  
  auc <- trapz(x,y)
  
  famy.vec[i] <- auc
}

#Create the output data frame consisting of the id and famy for each animal

output <- as.data.frame(cbind(id.vec, famy.vec))
output$famy.vec <- as.numeric(output$famy.vec)
output <- output%>% rename(id = id.vec,
                           famy.new = famy.vec)

#Create an additional output variable in which 0.3*lifespan is subtracted
#Create a data frame with each id and its lifespan, merge to the output, and create grail.fi as famy - 0.3*lifespan
max.doa <- input %>% group_by(id) %>% top_n(1, doa) %>% select(id, doa)
output <- output %>% left_join(max.doa) %>% mutate(grail.fi = (famy.new - 0.3*doa)) %>% select(id, famy.new, grail.fi)

#export the results as a csv file for further use
output.path <- "C:/Users/dlamming/Documents/Manuscripts in Preparation/QAMY paper/edited for second submission to Nature aging/Sample Data Out.csv"

write.csv(output,file = output.path, 
          row.names = FALSE)
